# Blockade of Interleukin-6 (IL-6) Signaling in Dedifferentiated Liposarcoma (DDLPS) Decreases Mouse Double Minute 2 (MDM2) Oncogenicity via Alternative Splicing

**DOI:** 10.1101/2024.02.21.581397

**Authors:** Abeba Zewdu, Danielle Braggio, Gonzalo Lopez, Kara Batte, Safiya Khurshid, Fernanda Costas de Faria, Hemant K. Bid, David Koller, Lucia Casadei, Katherine J. Ladner, David Wang, Valerie Grignol, O. Hans Iwenofu, Dawn Chandler, Denis C. Guttridge, Raphael E. Pollock

## Abstract

Effective therapies for retroperitoneal (RP) dedifferentiated liposarcoma (DDLPS) remain unavailable. Loco-regional recurrence occurs in >80% of cases; 5-year disease-specific survival is only 20%. DDLPS is especially prevalent in the retroperitoneum and abdomen; evaluation of the DDLPS microenvironment in these high-fat compartments appears pertinent. Adipose is a main supplier of interleukin-6 (IL6); excessive activation of IL6 signal transducer glycoprotein 130 (GP130) underlies the development of some diseases. The role of GP130 pathway activation remains unstudied in DDLPS, so we examined the role of microenvironment fat cell activation of the IL6/GP130 signaling cascade in DDLPS. All DDLPS tumors and cell lines studied expressed elevated levels of the GP130-encoding gene *IL6ST* and GP130 protein compared to normal tissue and cell line controls. IL6 increased DDLPS cell growth and migration, possibly through increased signal transducer and activator of transcription 1 (STAT1) and 3 (STAT3) activation, and upregulated mouse double minute 2 (MDM2). GP130 loss conveyed opposite effects; pharmacological blockade of GP130 by SC144 produced the MDM2 splice variant MDM2-ALT1, known to inhibit full length MDM2 (MDM2-FL). Although genomic *MDM2* amplification is pathognomonic for DDLPS, mechanisms driving MDM2 expression, regulation, and function beyond the MDM2:p53 negative feedback loop are poorly understood. Our findings suggest a novel preadipocyte DDLPS-promoting role due to IL6 release, via upregulation of DDLPS MDM2 expression. Pharmacological GP130 blockade reduced the IL6-induced increase in DDLPS MDM2 mRNA and protein levels, possibly through enhanced expression of MDM2-ALT1, a possibly targetable pathway with potential as future DDLPS patient therapy.

## Introduction

Dedifferentiated liposarcoma (DDLPS) is a high grade tumor of unknown etiology, characterized by wildtype *TP53* and genomic amplification of the p53 regulator *MDM2*. Surprisingly, targeted anti-MDM2 therapy displays modest impact with significant hematological toxicity (1,2), rendering radical surgery coupled with non-targeted toxic chemotherapies the mainstay treatment strategy, which often results in quality of life consequences (3,4).

These treatment approaches have not improved 5- and 10-year disease-free survival rates of 20% and 10%, respectively, which have stagnated since the 1970s (5–7). Although little is known about the DDLPS tumor microenvironment (TME), we identified a tumor-promoting paracrine loop linking DDLPS cells and tumor-associated macrophages (TAMs) (8). DDLPS cells were observed to release microRNAs (miR) 25-3p and 92a-3p in extracellular vesicles which then interacted with macrophage toll-like receptors 7 and 8, inducing elevated IL6 secretion and consequently enhancing DDLPS growth, migration, and invasion (8).

DDLPS proclivity to arise in the fat-rich retroperitoneum raises questions regarding how microenvironment adipose tissues might promote DDLPS progression. Adipose tissue is implicated as a key driver of development and progression of some cancers, in part by releasing pro-inflammatory cytokines such as IL6 (9–15). Interestingly, preadipocyte (PreAdip) cells generally secrete greater levels of IL6 compared to more differentiated adipocytes (16). IL6 binds to the GP130 receptor, subsequently inducing activation of disease-promoting downstream STAT1/STAT3, PI3K/AKT, and ERK/MAPK effector pathways (17–21). Although the impact of IL6 is incompletely understood, its activation of GP130 signaling participates in inflammation and development of some diseases (22,23). Moreover, GP130 pathway inhibition may convey potent anti-tumor effects (24), so we examined PreAdip cells as a possible source of IL6 and its impacts on DDLPS oncogenic behaviors via GP130 signaling.

Studies evaluating the impact of IL6 signaling on DDLPS MDM2 expression *per se* or *MDM2* splice variant production are lacking. Although full length *MDM2* is the most commonly expressed form(25), truncated variants produced by alternative mRNA splicing have been shown to be generated in response to genotoxic stress (26–28). MDM2-ALT1 splicing variant was found to be upregulated upon ultra-violet (UV) or *cis*-platinum treatment in osteosarcoma as well as breast and lung carcinomas bearing wild-type or deleted *TP53*, respectively, suggesting that MDM2-ALT1 production may be independent of *TP53* status (26). To our knowledge, no studies have evaluated the mechanisms driving alternative DDLPS *MDM2* splicing variant production. Given the genomic amplification of *MDM2* in DDLPS as well as the general lack of knowledge regarding DDLPS MDM2 regulation and alternative splicing, we evaluated the impact of IL6 signaling and blockade on full length and truncated *MDM2* expression.

## Materials and methods

### Reagents and drugs

SC144 was purchased from Sigma-Aldrich (cat# SML0763; St. Louis, MO, USA). Aliquots of SC144 were reconstituted in DMSO (cat# BP231-100; Fisher Bioreagents, Pittsburg, PA, USA) for *in vitro* studies and stored at −20°C until use. Recombinant human IL6 (cat# 206-IL-050) was purchased from R&D Systems (Minneapolis, MN, USA) and stored at −80°C.

### Cell culture

Human DDLPS cell lines, Lipo246, Lipo815, Lipo224, and Lipo863 were generated as previously described (29,30). LPS141 was attained from Dr. Jonathan Fletcher (Brigham & Women’s Hospital, Boston, MA, USA). Tumor cells were cultured in DMEM (Thermo-Fischer, Waltham, MA, USA) supplemented with 10% heat inactivated FBS (Gemini Bioproducts, Sacramento, CA, USA) and 1% penicillin/streptomycin (Life Technologies, Waltham, MA, USA). Immortalized PreAdip ( Lonza, Basel, Switzerland) and cultured in the appropriate media (SKBM-2 Bullet kit, Lonza, Basel, Switzerland: media supplemented with 0.5 mL human endothelial growth factor, 0.5 mL dexamethasone, 10 mL L-glutamine, 50 mL FBS, and 0.5 mL gentamycin) as described by supplier protocol. PreAdip cells were used as normal controls for *in vitro* assays. All cells were maintained at 37°C at 5% CO2 for the duration of the experiments. All cell lines tested negative for mycoplasma.

### Cell growth analysis

Cell growth endpoint analyses were performed using CellTiter96 Aqueous Non-Radioactive Cell Proliferation MTS Assay kit (Promega, Madison, WI, USA) per manufacturer specifications. To assess the impact of IL6 on DDLPS growth, tumor cells were seeded at a density of 2,000 cells per well in a 96-well plate in complete culture media The following day, cells were serum-starved by the removal of complete media and washing with PBS (Life Technologies, Waltham, MA, USA) prior to the addition of 100 μL plain DMEM to each well for 24h. On day 3, all media was replaced with fresh media containing plain DMEM with PBS (control) or 10 ng/mL IL6. After 96h, 20 μL of MTS was added to each well and cells were incubated at 37°C for 2h; absorbance was measured at 490 nm wavelength (Beckman Coulter, Indianapolis, IN, USA).

The effect of PreAdip-derived conditioned media (CM) on cancer cell growth was similarly evaluated. DDLPS cells were seeded at a density of 2,000 cells per well in a 96-well plate and 2.0×10^6^ PreAdip cells were seeded in a 10 cm dish in complete media. On day 2, the PreAdip media was removed, cells were washed twice with PBS, and fresh 5 mL DMEM supplemented with 0.5% bovine serum albumin (BSA, Amresco, OH, USA) was added to each plate. Also on day 2, the media was removed from the DDLPS wells, cells were washed with PBS and serum starved for 24h in plain DMEM. The following day, the PreAdip CM was collected and centrifuged (3000 rpm, 5 min, RT). Media was removed from the well containing DDLPS cells, and was replaced with 100 μL PreAdip-derived CM. Cells within the control group received 100 μL plain DMEM supplemented with 0.5% BSA. After 96h, endpoint assessment was performed using MTS.

The effect of IL6 or anti-IL6 monoclonal antibody MAB206 on the pro-growth impact of PreAdip CM on DDLPS cells was assessed using 2,000 cells per well. After adherence, cells were serum starved for 24h prior to the addition of plain DMEM supplemented with 0.5% BSA in the absence or presence of CM, 10 ng/mL IL6, 0.6 μg/mL MAB206, CM with 0.6 μg/mL MAB206, or CM with 10 ng/mL IL6. After 96h, cellular growth was measured using MTS reagent.

To assess the anti-DDLPS effects of SC144, cells were seeded at a density of 4,200 cells per well and allowed to adhere overnight. Treatment commenced the following day, and cellular growth was assessed 96h after treatment with DMSO (control), IL6 (10ng/mL), or increasing doses of SC144 (Sigma-Aldrich, St. Louis, MO, USA) in the presence of IL6 (10ng/mL) by MTS

### Gene knockdown

Cells seeded at a density of 1.75×10^5^ cells per well in a 6-well plate were transfected with 40 nM of pooled anti-IL6ST (cat# s7317, s7318, s7319; Thermo Fisher Scientific) or non-targeting Silencer Select siRNA (Thermo Fisher Scientific) in Opti-MEM media (Life Technologies) with Xtreme Gene transfected reagent (Sigma-Aldrich). The control cells were incubated only with transfection reagent and Opti-MEM.

### ELISA

Cells were seeded at a density of 1.75×10^5^ cells per well of a 6-well dish and, on the following day were washed once with PBS and media was replaced with fresh DMEM supplemented with 0.5% BSA. After 72h, supernatant was collected and centrifuged (3000 rpm, 5 min, RT) and Secreted IL6 was measured using the Quantikine® ELISA kit (R&D Systems), following manufacturer’s instructions.

### Complementary DNA synthesis and quantitative real-time PCR

Complement DNA (cDNA) synthesis was performed using TaqMan® Reverse Transcription Reagents (Life Technologies) and analyzed by quantitative real-time PCR (qRT-PCR) using TaqMan® Fast Advanced Master Mix ( Thermo Fisher Scientific) and MDM2 TaqMan probe (Hs01066930_m1) or IL6ST TaqMan probe (Hs00174360_m1) (Life Technologies). Relative gene expression levels were normalized to *GAPDH* expression (TaqMan probe cat# Hs02758991_g1).

cDNA was also generated using iScript™ Reverse Transcription Supermix (Bio-Rad Laboratories, Hercules, CA, USA) per manufacturer’s specifications and assessed by qRT-PCR using SYBR® Green PCR Master Mix (Thermo Fisher Scientific). Primer sequences are listed below, relative gene expression was normalized to *GAPDH*.

IL6 (F: CACTCACCTCTTCAGAACCAAT; R: GCTGCTTTCACACATGTT ACTC)

GAPDH (F: GAAGGTGAAGGTCGGAGTC; R: GAAGATGGTGATGGGATTTC)

PUMA (F: ACGACCTCAACGCACAGTACGA; R: GTAAGGGCAGGAGTCCCATGATGA)

The StepOnePlus Real-Time PCR System (ThermoFisher Scientific) was used to execute the quantitative real-time PCR reactions.

### Western blot

Protein extract was isolated from cells using 1x Cell Signaling Lysis Buffer (Cell Signaling, Danvers, MA, USA), supplemented with 1x Halt™ Protease Inhibitor Cocktail EDTA-free (Thermo Fisher Scientific).

For western blot analysis, 25 μg protein were subjected to electrophoretic run using a 4–15% Mini-PROTEAN® TGX™ Precast Protein Gels (Bio-Rad Laboratories), transferred to PVDF membranes using the Trans-Blot® Turbo™ Midi PVDF Transfer Packs (Bio-Rad Laboratories) and dried for at least 1h. Membranes were incubated overnight at 4°C with the antibodies listed below. Primary antibody mixtures were removed, and membranes were washed and then incubated with secondary donkey anti-rabbit or donkey anti-mouse (IRDye 800CW) and/or donkey anti-goat, donkey anti-mouse, or donkey anti-rabbit (IRDye 680RD) (Li-Cor Biosciences, Lincoln, NE, USA) antibody for 1h. Membranes were washed and dried. Imaging and protein expression analyses were performed with the Odyssey CLx imaged (Li-Cor). Antibodies: STAT1 (cat# 9176), pSTAT1 (Tyr701; cat# 9167), ERK (cat # 4696S), pERK1/2 (Thr202, Tyr204; cat# 4370S), AKT (cat# 4691), pAKT (Ser473; cat# 12694), GP130 (cat# 3732), STAT3 (cat# 4904), pSTAT3 (Tyr705; cat# 4113), and pSTAT5 (Tyr694; cat# C11C5), from Cell Signaling; MDM2 (cat# MA1-113) and STAT5 (cat# 13-3600), from Thermo Fischer Scientific; GAPDH (cat# sc-20357), from Santa Cruz Biotechnology (Dallas, TX, USA).

### Migration assay

Cellular migration was assessed using 8μm trans-well ThinCerts™ migration chambers (Geiner Bio-One, Kremsmünster, Austria). DDLPS cells were seeded to 60% confluence in a 10 cm dish in complete culture media. The following day, cells were treated with PBS, 10 ng/mL IL6, DMSO, or 5 μM SC144 plus 10 ng/mL IL6 for 24h. The next day, treated cells were quantified and 300,000 viable cells were seeded into the upper chamber of the migration chamber in 200 μL plain DMEM. Antibiotic-free DMEM with 5% FBS served as chemoattractant. Endpoint staining was performed by fixing cells in 0.5% crystal violet in ethanol (Fisher Scientific) for 1h with gentle rotation, then washing the migration chambers gently three times with double-distilled water. Chambers were imaged after drying overnight or longer.

### Indirect and direct co-culture analyses

The pro-DDLPS impact of culturing PreAdip cells with DDLPS cell lines was determined by both direct and indirect co-culture. Co-culture assays in which PreAdip cells were cultured directly with DDLPS cells were performed using red fluorescent protein (RFP)-tagged DDLPS cells. 2,000 PreAdip cells were cultured into the basement well of a 96-well plate in 100 μL complete DMEM. The following day, DMEM + 0.5% BSA was added to each PreAdip well. 2,000 RFP-tagged DDLPS cells were re-suspended in DMEM + 0.5% BSA and added to each well housing PreAdip cells (co-culture). Cells were observed for 5 days and viable cells were quantified using the IncuCyte Zoom live-imaging fluorescent microscope system. The total fluorescent signal per each well indicates the number of viable, fluorescently-tagged DDLPS cells in that respective well.

Indirect co-culture experiments were performed by culturing DDLPS cells with PreAdip cells in 6-well Boyden 0.4 μm pores chambers (Corning Costar, Corning, NY, USA)). These experiments were performed as follows: On day 1, tumor cells were plated at a density of 2.0 x 10^5^ cells/well into the basement well of a 6-well plate in 2 mL complete media (DMEM + 10% FBS + 1%P/S). Also on day 1, 10^5^ PreAdip cells were seeded into the insert bearing the perforated membrane, and placed into a second 6-well plate containing complete PreAdip media. The following day, the wells with the DDLPS cells and the inserts with PreAdip cells were washed prior to the addition of plain DMEM supplemented with 0.5% BSA for 24h. On day 3, all media was aspirated and replaced with fresh DMEM plus 0.5% BSA, and the inserts bearing the PreAdip cells were placed into the wells with the adherent DDLPS cells. Cancer cells cultured in the absence of PreAdip cells in plain DMEM supplemented with 0.5% BSA served as the monoculture control for each respective DDLPS cell line. Endpoint analysis of DDLPS proliferation was performed at day 5, by quantifying viable cells count using trypan blue exclusion.

### Cell viability

DDLPS cell viability in response to SC144 treatment was assessed using RFP-tagged DDLPS cells using the IncuCyte Zoom system detailed above. In short, 4,200 cells per well were plated and treated with increasing doses of SC144 in the presence of IL6 (10 ng/mL) and were monitored by live-imaging fluorescent microscopy over the course of 96h. Quantification of RFP-fluorescing cells indicated live DDLPS cells.

### Flow cytometry

Apoptosis was assessed using the Apoptosis Detection Kit (BD Pharmagen, San Diego, CA, USA). Following supplier instructions, 10^6^ cells/mL of cells treated with DMSO, IL6 (10ng/mL, SC144 (5uM), or IL6 plus SC144 for 96h were stained with 5 uL Annexin V-FITC and 5 μL Propidium iodide (PI, Sigma-Aldrich, St. Louis, MO, USA) and analyzed by flow cytometry (LSR II, BD Pharmagen, San Diego, CA, USA).

### Nested PCR for MDM2 splicing variant assessment

RNA was extracted from Lipo246 and Lipo815 cell lines and Reverse transcription (RT) reactions were carried out using Transcriptor RT enzyme (Roche Diagnostics, Basel, Switzerland). Nested PCR was performed using a 2-step PCR reaction as previously described(31).

External Sn: 5’-CTGGGGAGTCTTGAGGGACC-3’

External ASn: 5’-CAGGTTGTCTAAATTCCTAG-3’

Internal Sn: 5’-CGCGAAAACCCCGGGCAGGCAAATGTGCA-3’

Internal ASn: 5’-CTCTTATAGACAGGTCAACTAG-3’

### In vivo xenograft model

Six week old female athymic nude (NCr-nu/nu) mice, outbred, were acquired from the athymic nude mouse colony maintained by the Target Validation Shared Resource at the Ohio State University; original breeders (strain #553 and #554) for the colony were received from the NCI Frederick facility, and were injected subcutaneously with 10^6^ Lipo246 cells into the flank of 20 mice. Upon reaching 0.5 cm^3^ tumor size, the 15 mice that displayed tumor formation were randomized into two arms and daily treated via intraperitoneal injections: Vehicle (0.9% NaCl/40% propylene glycol/59.1% ultra-pure distilled water; 8 mice), or SC144 (10 mg/kg diluted in vehicle solution; 7 mice). Mice were weighed and tumors were measured twice weekly. Mice were euthanized once tumors in the control group grew to ∼1.5 cm^3^. Final tumor volumes and weights were measured; tumor volume was calculated using the formula (L x (W^2^)/2.

### Study approval

Approval for the proper handling of mice was obtained from The Ohio State University Office of Responsible Research Practices in adherence with the Institutional Animal Care and Use Committee protocols (approval #2014A00000085) in accordance with The Ohio State University’s Animal Welfare Assurance (#A3261-01). DDLPS tumor samples were provided by patients of the James Cancer Center of The Ohio State University Wexner Medical Center. Written, informed consent to participate in the study was obtained from each donor, and approval for the handling of patient samples was attained from The Ohio State University Internal Review Board (approval #2014C0028).

### Immunohistochemistry analysis

Antibodies GP130 (cat# ab170257) and IL6 (cat# ab6672), Abcam, Cambridge, United Kingdom, were used for immunohistochemistry (IHC) assessment at a dilution of 1:250. IHC staining was conducted at Nationwide Children’s Hospital (Columbus, OH, USA) and the Ohio State University College of Veterinary Medicine, and analysis was performed at the Polaris Innovation Center (The Ohio State University, Columbus, OH, USA).

### Statistics

Mean ± SEM values were generated for all cell-based assays, and statistical analyses and EC_50_ values were computed using GraphPad Prism version 6.00 (GraphPad Software, La Jolla California USA, www.graphpad.com). All Student *t* tests performed were unpaired and two-sided, with Welch’s correction applied to the analysis of patient *IL6ST* mRNA expression. Analyses across multiple cell lines were performed by one-way ANOVA.

During the investigation of the *in vivo* impact of SC144 on tumor growth, mean ± SEM was calculated for each treatment group, and end-point analyses determined using unpaired two-sided *t* test was utilized to determine variances. The average tumor volume (mm^3^) and weight (g) for each study arm was measured and recorded. Significance was set at \**p* ≤ 0.05 for all *in vitro* and *in vivo* experimentation.

## Results

### GP130 is overexpressed in DDLPS and requires an extracellular cytokine signal for activation

To determine if GP130 signaling facilitated DDLPS progression, expression of GP130 in DDLPS tumors (S1 Table) and cell lines (S2 Table) was examined. *IL6ST* (the gene that encodes GP130 protein) and GP130 measurements revealed consistently high DDLPS tumor and cell line expression (Figs 1A-D), suggesting possible relevance to DDLPS development and/or progression.

**Fig 1.**
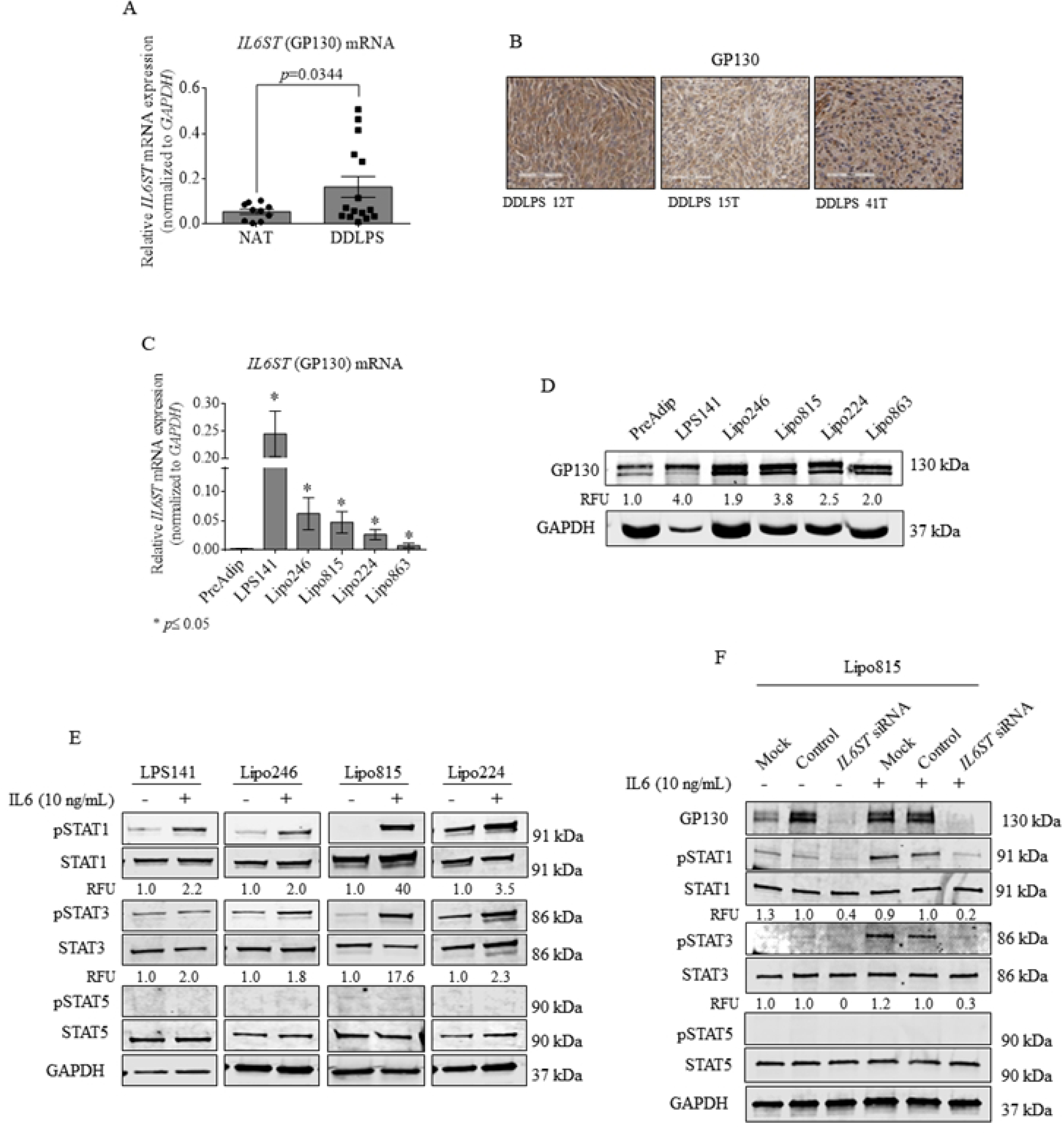
GP130 is elevated in DDLPS; receptor activation requires exogenous IL6. (A) *IL6ST* expression in DDLPS patient tumors (n= 15) versus normal adjacent tissue (NAT; n= 10). (B) Representative 20x magnification of DDLPS patient tumor GP130 IHC staining. (C) DDLPS and normal PreAdip control cell *IL6ST* expression. (D) GP130 protein levels in established DDLPS cell lines. (E) STAT1, STAT3 and STAT5 protein expression in DDLPS cell lines after IL6 cytokine treatment (10 ng/mL, 20 min) and (F) after *IL6ST* KD (48h) in Lipo815. STAT5 activation, which is regulated by IL2 signaling, remained unaffected by GP130 activation or loss. KD, knockdown.

GP130 signal transduction activates downstream STAT1 and STAT3 effector proteins (17–19). STAT1 and STAT3 were inactive or marginally activated in DDLPS cells, but showed enhanced activation with exogenous IL6 (Fig 1E). Loss of DDLPS GP130 severely reduced active STAT1 and STAT3 in the presence of IL6, suggesting that their activation by IL6 occurs in a GP130-dependent manner (Fig 1F). AKT remained inactivated in DDLPS cells, and the addition of IL6 did not impact AKT or ERK1/2 activation (S1 and S2 Figs). Moreover, DDLPS cells express constitutively active ERK1/2 independent of IL6 GP130 activation (S1 and S2 Figs).

### GP130 activation and loss conversely affect DDLPS protumorigenic properties

IL6-mediated activation of STAT1 and STAT3 by inducing malignant proliferation, dissemination, and attenuated therapeutic disease responses (20,21,32–37). However, STAT1 and STAT3 roles in DDLPS remain poorly understood. We assessed the impact of GP130 activation on DDLPS growth and migration. DDLPS cells cultured in the presence of IL6 demonstrated increased growth after 96h (Lipo246: 127% ± 2.23, *p*≤0.001; Lipo815: 117% ± 7.13, *p*≤0.01; Fig 2A) and elevated migration (Fig 2B). Conversely, *IL6ST* siRNA knockdown (KD) decreased cellular growth at 96h (Lipo246: 76.11% ± 1.15, *p*=0.0023; Lipo815: 77.82% ± 4.382, *p*=0.0023; Fig 2C). PreAdip cell growth remained unaffected by the activation or loss of GP130 (Figs 2A and 2C).

**Fig 2.**
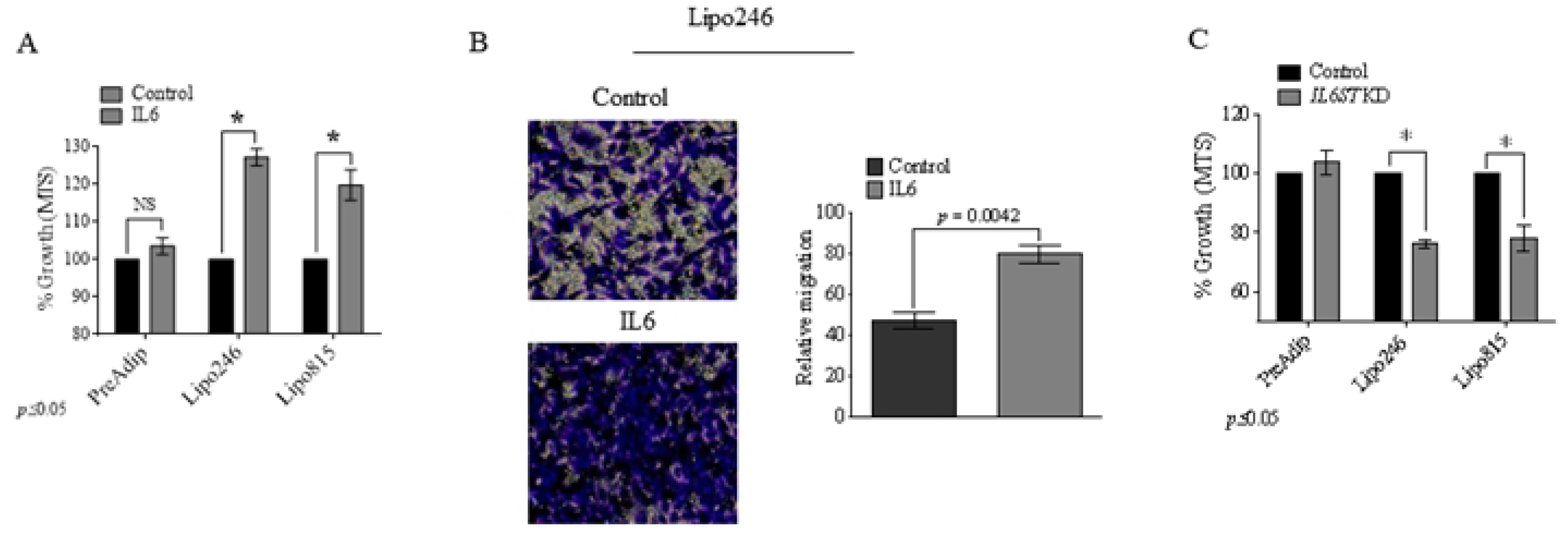
GP130 activation promotes DDLPS oncogenic phenotype. (A) MTS endpoint analysis of DDLPS and preadipocyte growth with IL6 (10 ng/mL, 96h). (B) DDLPS Lipo246 migration after treatment with IL6 (10 ng/mL, 24h; *p*=0.0042). (C) MTS assessment of preadipocyte, Lipo246 (*p*=0.0023) and Lipo815 (*p*=0.0023) cells after GP130 *KD* (96h).

### Preadipocytes may act as an *in situ* source of IL6 for DDLPS tumors

Patient sample immunohistochemistry (IHC) was performed to detect IL6 within DDLPS TME, revealing IL6 expression in DDLPS tumors (Fig 3A). Consistent with others(16), we observed an elevated expression and secretion of IL6 by PreAdip and DDLPS cells, albeit DDLPS expressed substantially lower IL6 levels (Fig 3B). DDLPS cells cultured directly or indirectly with PreAdip cells displayed markedly increased proliferation compared to DDLPS monoculture cells (Figs 3C and D). DDLPS cells cultured in PreAdip-derived conditioned media (CM) demonstrated similar growth increases that were not further augmented by exogenous IL6 (Fig 3E). Anti-IL6 neutralizing antibody (MAB206) reduced the DDLPS-promoting effect of PreAdip cells with simultaneous reduction of DDLPS GP130 activation (S3 and S4 Figs). However, treatment with MAB206 did not fully inhibit the pro-DDLPS growth effects of PreAdip, suggesting that these normal cells may also promote DDLPS growth by mechanisms other than IL6 release.

**Fig 3.**
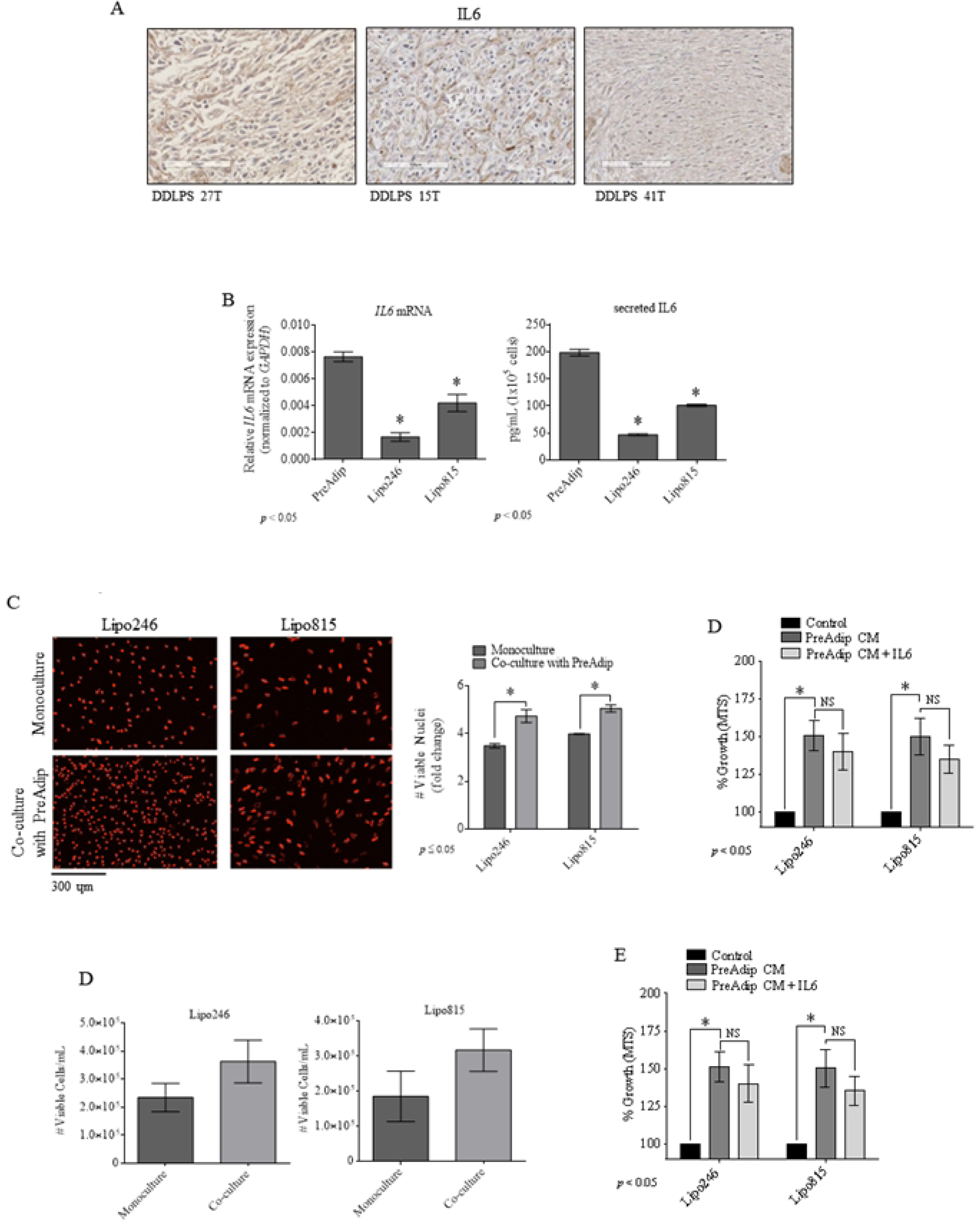
Preadipocytes may promote DDLPS growth *in situ*. (A) IHC staining of IL6 of DDLPS tumor tissue. (B) IL6 mRNA (*left*) and secreted protein levels (*right*) in PreAdip and DDLPS cells. (C) RFP-tagged DDLPS cells cultured alone or co-cultured with PreAdip cells (untagged). (D) Cellular viability of DDLPS cells indirectly co-cultured with PreAdip cells (day 5). (E) DDLPS cell growth with PreAdip-derived CM (96h).

### GP130 signaling increases DDLPS MDM2 expression

Given the consistent overexpression of MDM2 in DDLPS tumors, we considered possible GP130 impacts on MDM2. DDLPS cells were cultured under serum-deprived conditions followed by IL6 treatment. Lipo246 and Lipo815 DDLPS cells demonstrated 3- and 10-fold rises in *MDM2* transcripts, respectively, and increased MDM2 protein levels (Fig 4A), whereas siRNA-mediated KD of *IL6ST* in Lipo246 cells resulted in decreased in MDM2 expression (Fig 4B).

**Fig 4.**
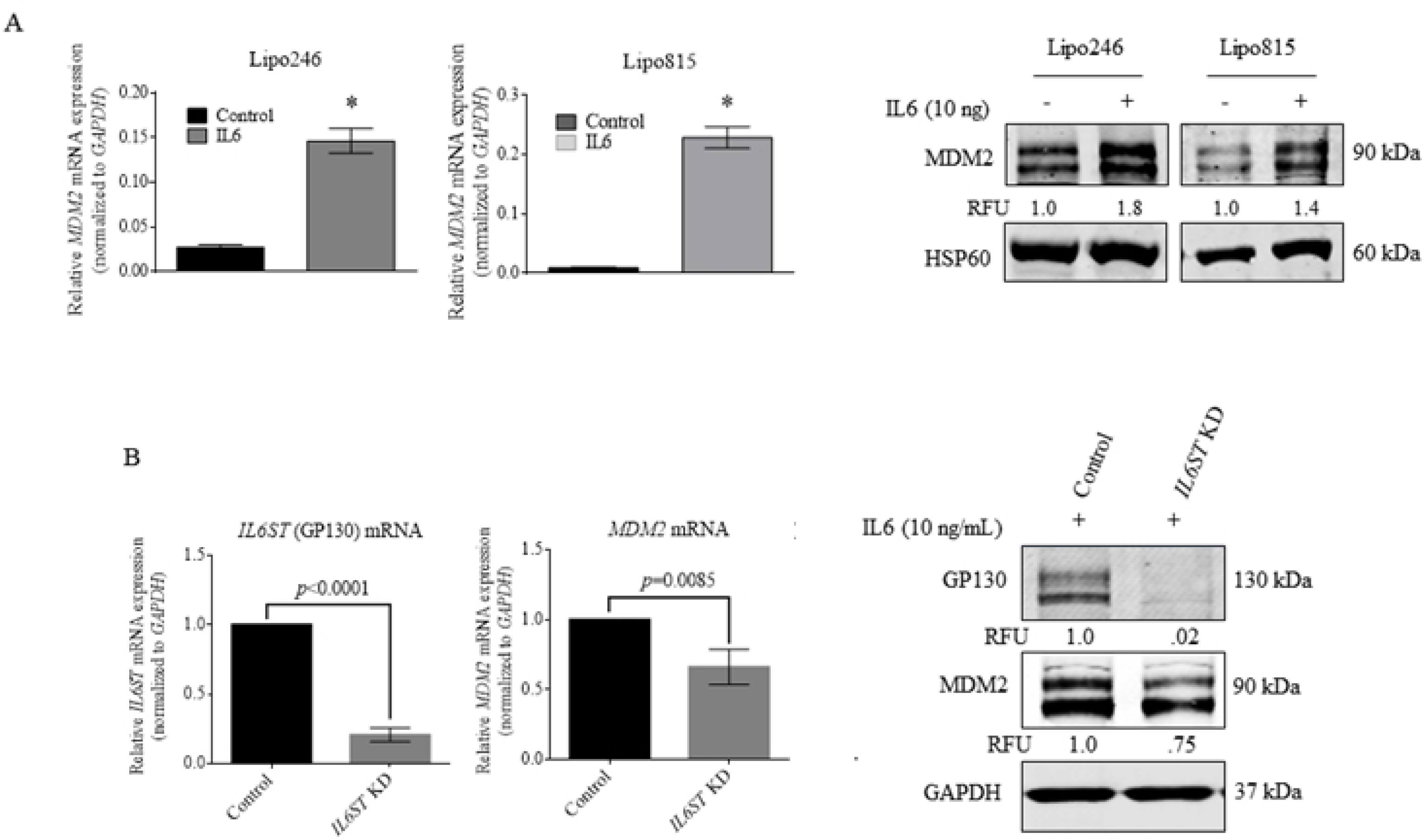
GP130 activation and loss in DDLPS affects MDM2 levels. (A) *MDM2* mRNA expression (*top*; *\*p*≤0.05) and protein levels (*bottom*) in DDLPS after 96h IL6 treatment. (B) MDM2 and GP130 expression in Lipo246 cells following *IL6ST* KD.

### GP130 blockade by SC144 reduces DDLPS growth *in vitro* and *in vivo*

Our findings suggested potentially protumorigenic GP130 roles in DDLPS, so we investigated GP130 as a possible therapeutic target. MTS analysis of DDLPS treatment with anti-GP130 agent SC144 reduced *in vitro* DDLPS growth by 96h with a dose as low as 0.625 μM SC144 (Fig 5A), whereas PreAdip cells demonstrated treatment tolerance (EC_50_=6.332 μM SC144; S3 Table). Similarly, SC144 GP130 blockade inhibited Lipo246 and Lipo815 DDLPS cell proliferation, as indicated by reduced numbers of viable, red fluorescent protein (RFP)-tagged DDLPS cells (Fig 5B). SC144 treatment also abrogated DDLPS migration (Fig 5C) and induced approximately 5-and 3-fold increased apoptosis in Lipo246 and Lipo815 DDLPS cells, respectively (Fig 5D). Pharmacologic GP130 blockade induced *TP53* expression with upregulation of the proapoptotic gene *PUMA* (Fig 5E).

**Fig 5.**
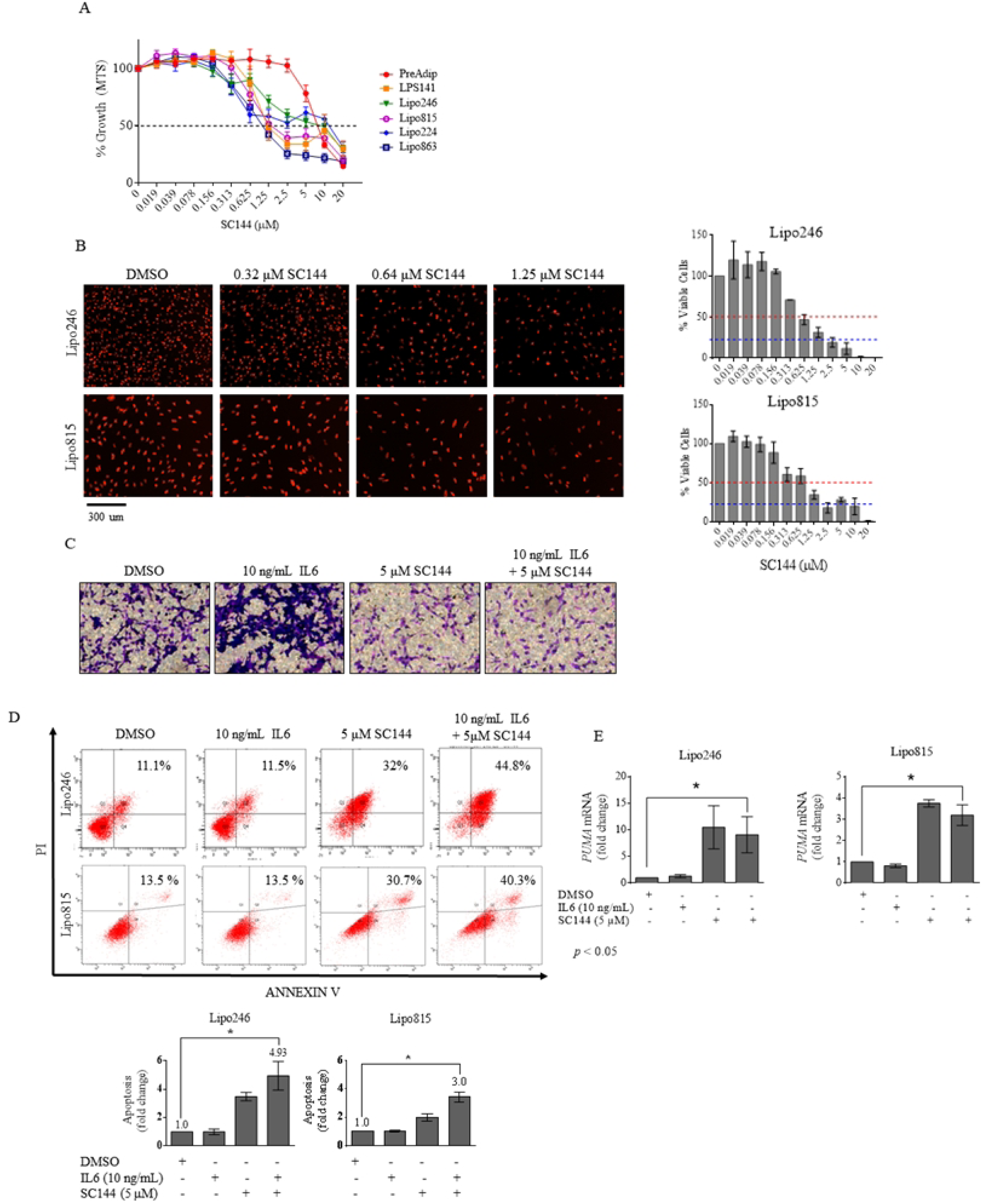
SC144 reduces DDLPS growth *in vitro*. MTS endpoint analysis of (A) cell growth and (B) proliferation of DDLPS and preadipocyte cells cultured in increasing doses of SC144 for 96h. (C) Migration evaluation of Lipo246 DDLPS cells treated with SC144 (5 μM, 24h). (D) Apoptosis analysis of Lipo246 and Lipo815 after SC144 (5 μM, 96h) treatment. (E) Proapoptotic gene *PUMA* expression after Lipo815 and Lipo246 tumor cell treatment with SC144. Tumor volume (F) and tumor weight (G) of mice bearing DDLPS Lipo246 tumors treated with SC144 vs vehicle control (*p*=0.0273). *\*p*≤0.05.

Our Lipo246 xenograft mouse model (29,30) was utilized to evaluate SC144 as an *in vivo* anti-DDLPS agent. Lipo246 cells were injected subcutaneously into the flank of athymic nude mice; treatment commenced once tumors reached 0.5 cm^3^. Mice bearing tumors were treated with vehicle or 10mg/kg SC144 via daily intraperitoneal injections. At termination, treatment with SC144 had significantly delayed tumor growth (1315.6 ± 329.1 mm^3^ v 2704.9 ± 435.4 mm^3^, *p*=0.0273; Fig 6A) and decreased tumor weight by ∼33% (1.58 ± 0.23g v 1g ± 0.24g; Fig 6B) versus controls.

**Fig 6.**
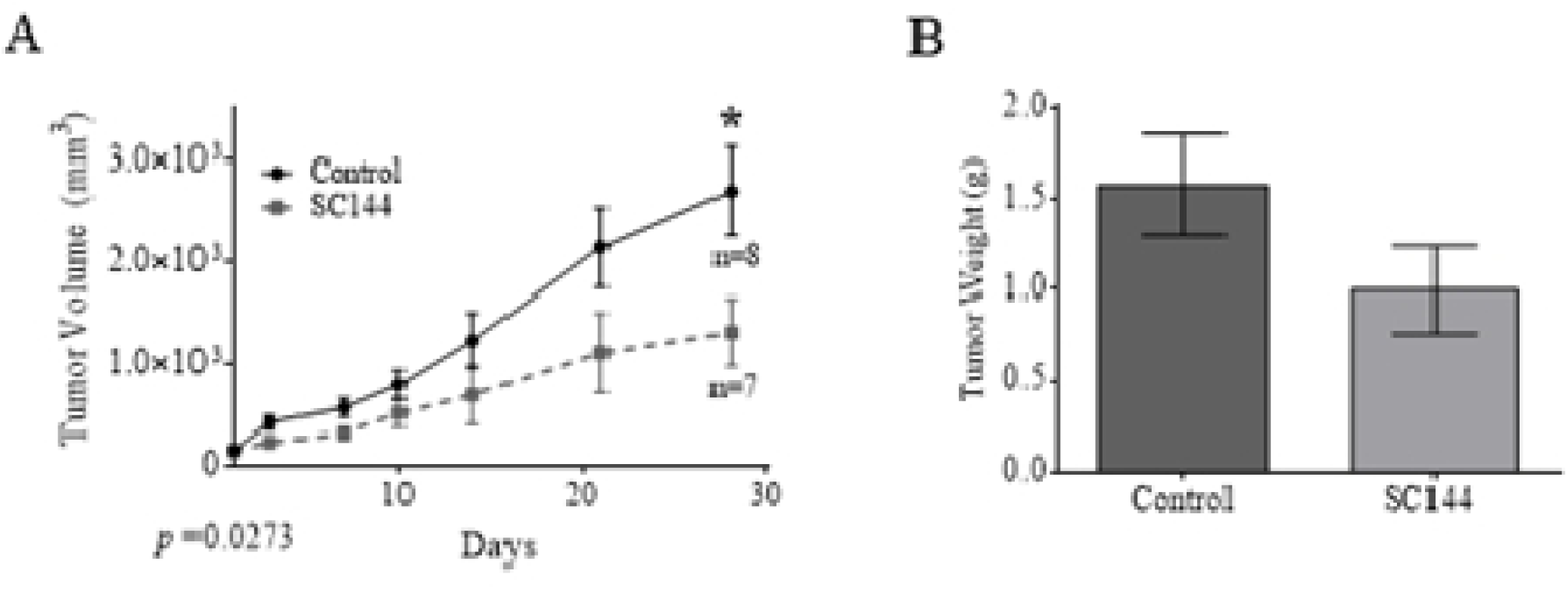
SC144-mediated GP130 blockade impairs DDLPS growth *in vivo*. Tumor volume (A) and tumor weight (B) of mice bearing DDLPS Lipo246 tumors treated with SC144 vs vehicle control (*p*=0.0273). *\*p*≤0.05.

### SC144 has GP130-specific effects on STAT1 and STAT3 activation, and reduces DDLPS MDM2 expression

Considering the GP130-specific effects of SC144 (24,38), we asked if SC144 would affect DDLPS GP130 signaling, perhaps via STAT1 and STAT3 activation. Interestingly, we observed a shift in GP130 expression from the higher to the lower molecular weight band, suggesting protein deglycosylation consistent with possible GP130 inactivation (Fig 7A) (24). Levels of pSTAT1 and pSTAT3 decreased concurrently with GP130 blockade (Fig 7B). Similarly, nuclear expression of activated STAT1 and STAT3 also decreased (Fig 7C).

**Fig 7.**
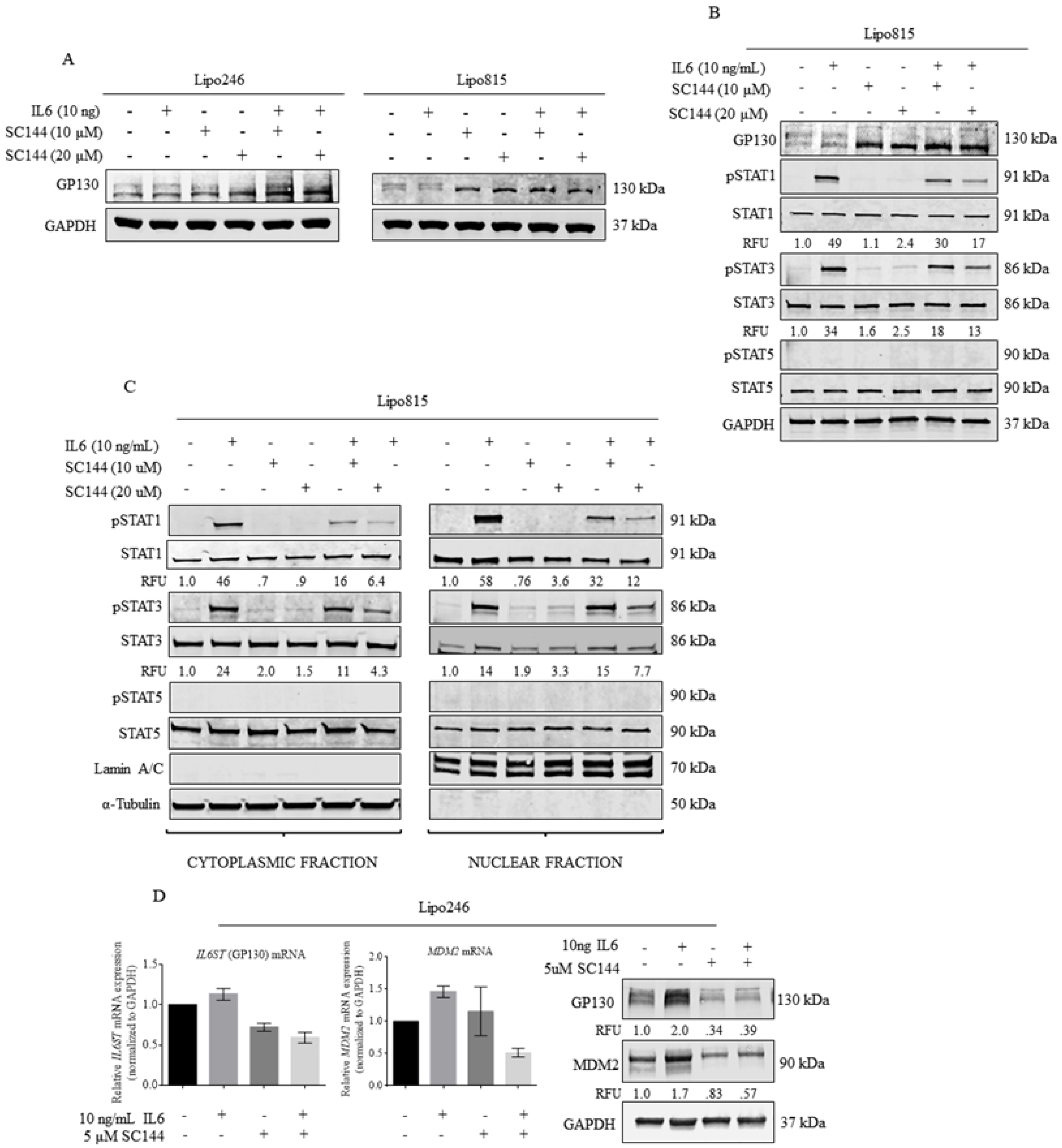
SC144-mediated GP130 inhibition impairs STAT1 and STAT3 activation, and reduces DDLPS GP130 and MDM2 expression *in vitro*. (A) GP130 protein expression in Lipo246 and Lipo815 cells after SC144 treatment. (B) Whole-cell and (C) nuclear and cytoplasmic STAT1 and STAT3 protein levels in Lipo815 cells treated with IL6 and SC144. (D) GP130 and MDM2 expression in Lipo246 cells treated with IL6 and/or SC144 (5 μM SC144 -/+ 10 ng/mL IL6, 48h).

To assess the impact of GP130 blockade on MDM2 expression, Lipo246 DDLPS cells were treated for 48h with SC144 with or without IL6, after which GP130 and MDM2 levels were measured. Treatment with SC144 significantly decreased expression of MDM2 mRNA and protein with concurrently decreased GP130 levels (Fig 7D), indicating possible GP130 signaling in MDM2 regulation. As observed *in vitro*, *IL6ST* and *MDM2* levels decreased in tumors of mice receiving SC144 treatment versus controls (Fig 8A).

**Fig 8.**
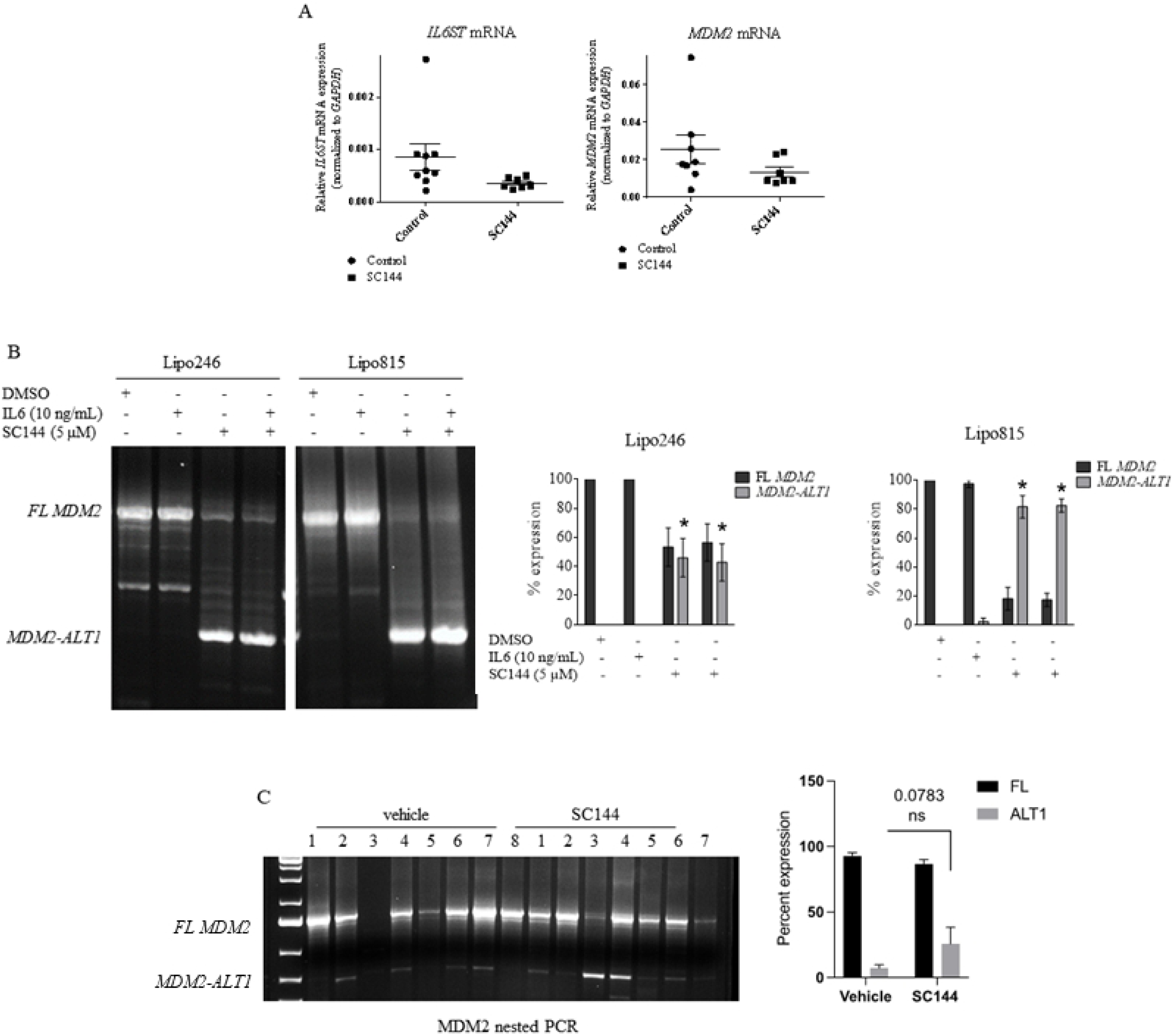
GP130 signaling blockade decreases MDM2 expression and upregulates MDM2-ALT1 expression *in vivo*. (A) *IL6ST* and *MDM2* mRNA expression with *in vivo* SC144 (10 mg/kg injected IP) treatment of Lipo246 tumor-bearing mice on day 28. Semi-quantitative expression analysis of *MDM2* isoforms *MDM2-*full length and *MDM2-ALT1* in DDLPS cells (B) and Lipo246 tumor-bearing mice (C) treated with SC144.

### GP130 blockade upregulates production of *MDM2-ALT1*, leading to DDLPS p53 reactivation

Full length MDM2 mRNA comprises a longer mRNA script than its truncated MDM2-ALT1 splice form, which is known to inhibit full length MDM2 (Supplemental Figure S5). Little is known about DDLPS alternatively spliced *MDM2* isoforms, so GP130 activation and blockade of *MDM2* splice variant expression was assessed by semi-quantitative PCR. Addition of IL6 increased *MDM2-FL* gene levels in Lipo246 and Lipo815 cells whereas GP130 blockade by SC144 decreased *MDM2-FL* expression while concurrently increasing *MDM2-ALT1* levels (Fig 8B). SC144 treatment similarly elevated *MDM2-ALT1* expression *in vivo* (Fig 8C). These findings suggest that DDLPS GP130 blockade via SC144 increases *MDM2-ALT1* production, leading to sequestration of full-length MDM2 and subsequent release of p53 from MDM2 regulatory mechanisms (Fig 9).

**Fig 9.**
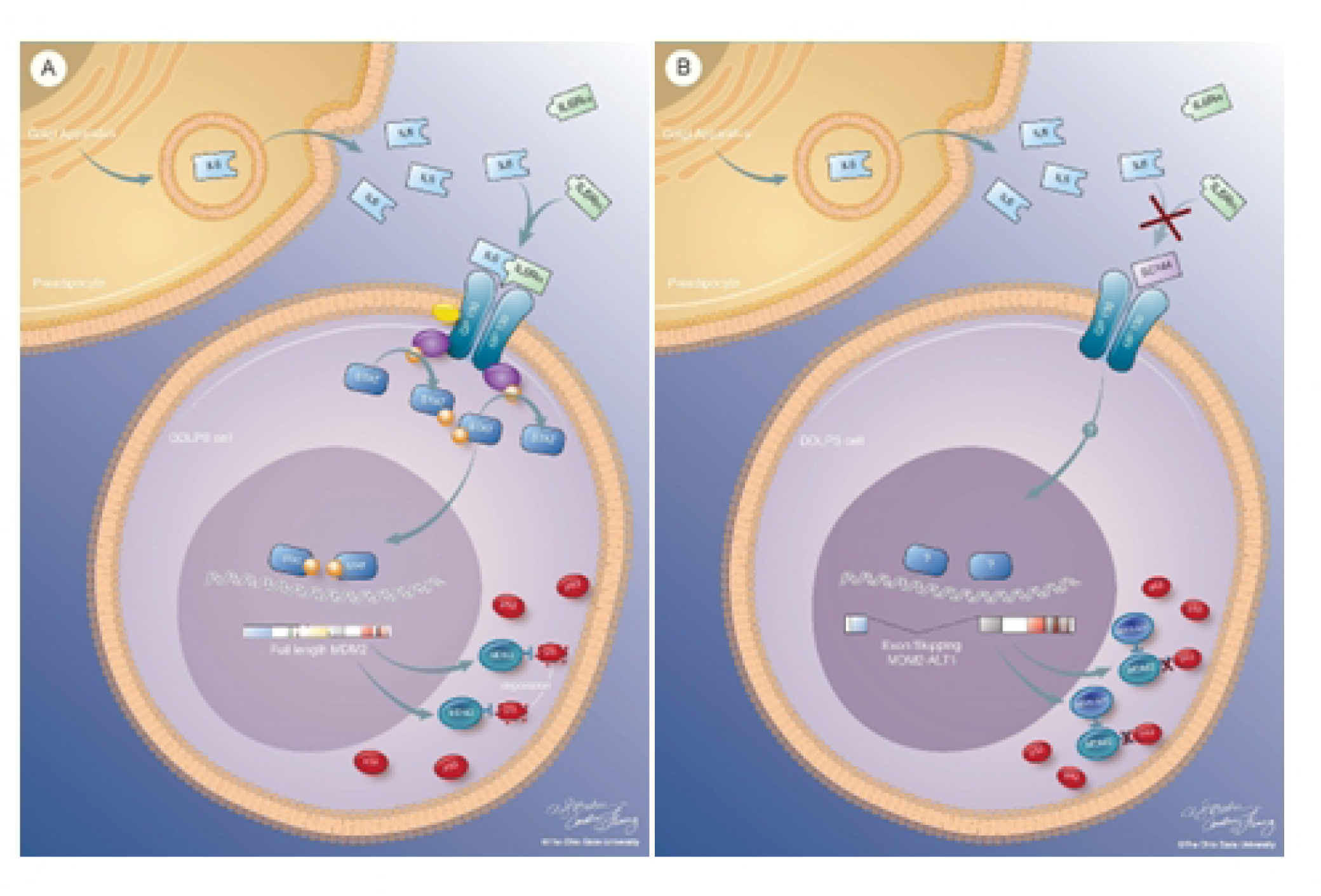
Proposed mechanism of GP130 signaling-mediated upregulation of *MDM2* and oncogene suppression via SC144-induced expression of MDM2-ALT1. Proposed mechanism of how IL6-mediated DDLPS GP130 activation leads to downstream upregulation of MDM2 (left) and SC144-mediated GP130 inhibition, resulting in MDM2-ALT1 production and decreased MDM2-FL expression (right).

## Discussion

To the best of our knowledge, we are the first to show a tumor-promoting paracrine loop in which DDLPS cells release microRNAs 25-3p and 92a-3p in extracellular vesicles which then interact with toll-like receptors 7 and 8 in macrophages, inducing elevated IL6 secretion. Consequently, we evaluated GP130 signaling in DDLPS, finding that IL6-rich TAM-derived conditioned media treatment enhances DDLPS growth, migration, and invasion (8). Here we demonstrate that PreAdip cells also release elevated levels of IL6, perhaps also acting to promote DDLPS. Signaling facilitated by IL6 is mediated by GP130, which is elevated in DDLPS patient samples and cell lines. Moreover, activation of GP130 by IL6 enhanced the oncogenic phenotype of DDLPS (i.e., increased growth and migration), perhaps by activating of downstream STAT1 and STAT3 effector proteins. Based on these findings, we then demonstrated that GP130 activation by IL6 increases pSTAT1 and pSTAT3 levels, leading to increased MDM2 expression.

Detection of elevated DDLPS *MDM2* copy number is a basis for disease diagnosis; however, mechanisms regulating its oncogenic expression are poorly understood. Our studies show that IL6 activation of GP130 results in elevated MDM2 expression whereas *IL6ST* KD results in reduced MDM2 levels, suggesting that DDLPS MDM2 expression is partially regulated by GP130 signaling. Upregulation of DDLPS MDM2 by GP130 signal transduction may be facilitated by STAT1 and STAT3 transcription factors, as seen in colorectal cancer (37) where GP130 activation by IL6 cytokine family member leukemia inhibitory factor led to increased MDM2 levels in a STAT3-dependent fashion(37). In CRCs harboring wildtype p53, this increase in MDM2 expression resulted in decreased apoptosis and increased chemoresistance. The MDM2 role in driving cancer drug resistance could underly the minimal DDLPS chemotherapeutic responses (39–43).

Synthetic anti-GP130 drugs have shown promising results. Xu *et al*. showed ovarian cancer tumor regression *in vivo* with the treatment of SC144 (24). SC144 treatment led to deglycosylation and subsequent phosphorylation and internalization of GP130, drastically reducing the level of surface-bound protein and IL6-induced GP130 signaling neutralization (24). Xiao similarly observed efficacy during the use of Bazedoxifene, an FDA-approved agent with known anti-IL6:GP130 axis activity currently used to prevent postmenopausal osteoporosis (44).

Genotoxic stress induces production of alternative MDM2 variants and modulates the MDM2:p53 axis, thereby affecting p53 activity (26,28,45). Jacob *et al.* demonstrated that alternative splice product MDM2-ALT1 was able to bind MDM2-FL and MDMX, the conventional MDM2 binding partner involved in p53 negative regulation(28). As MDM2-ALT1 lacks the p53-specific binding site but maintains the RING domain required for MDM2 dimerization, it is believed to physically impair MDM2-FL activity via direct binding (28). Cytoplasmic binding of MDM2-ALT1 to MDM2 rendered MDM2 inert, which resulted in p53 stabilization, the subsequent upregulation of downstream p53 target p21, and an arrest in cell cycle progression. We similarly observed substantial increases in MDM2-ALT1 and *TP53* levels with markedly elevated apoptotic levels and PUMA expression upon DDLPS treatment with SC144. These data suggest that p53 reactivation in response to GP130 blockade-induced MDM2-ALT1 production may impair DDLPS progression via apoptosis induction. Furthermore, the observed decreased MDM2-FL transcript and elevated MDM2-ALT1 expression upon GP130 blockade suggest that DDLPS MDM2 is negatively regulated on the genomic level as well as post-translationally.

Understanding the potential impact of TME non-cancer cells on DDLPS is critical, especially given the functional role of stromal cells in promoting cancer progression(46,47). Here, we have established a novel PreAdip:DDLPS paracrine loop in which PreAdip-derived IL6 enhances DDLPS growth and oncogenic phenotype, potentially via activation of downstream STAT1 and STAT3 effector proteins. Co-culture with DDLPS cells may also modify PreAdip biology. Others have described the transition of normal cells into cancer-promoting bodies in the presence of cancer cells(46,47). In a recent publication by Lee *et al.*, cancer-associated adipocytes were shown to undergo dedifferentiation and express higher levels of IL6 when co-cultured with breast cancer cells. These modified fat cells were subsequently able to promote cancer cell progression(47). Therefore, genetic, proteomic, and morphological analyses comparing cancer-associated cells with their normal cell counterparts could be considered to better understand the PreAdip:DDLPS cell relationship and further define the tumor-promoting functions of PreAdip cells within the TME. Results of these studies will hopefully enable novel treatment strategies based on better understanding of the heterogeneous DDLPS tumor microenvironment.

## Acknowledgement

We thank the Target Validation Shared Resource at the Ohio State University Comprehensive Cancer Center for providing the athymic nude mice used in preclinical studies. The content is solely the responsibility of the authors and does not necessarily represent the official views of the National Institutes of Health or the United States of America Department of Defense. The authors would also like to thank Dr. Jonathan Fletcher for his generous donation of the LPS141 DDLPS cell line as well as the Columbus Nationwide Children’s Hospital for its contribution to this work.

## Supporting information

**S1 Table.** Patient tissue samples and associated patient history.

**S2 Table.** Cell lines derived from DDLPS tumors.

**S3 Table.** EC_50_ values of normal and DDLPS cell lines determined by MTS after 96h treatment with SC144.

**S1 Fig. ERK1/2 is constitutively active in DDLPS.** AKT and ERK1/2 protein expression in serum-starved DDLPS cells (24h).

**S2 Fig. The addition of IL6 to serum-starved DDLPS cells did not alter AKT activation or pERK1/2 expression.** AKT and ERK1/2 protein expression in Lipo246 and Lipo815 cells after treatment with IL6 (10 ng/mL, 20 min).

**S3 Fig. IL6 neutralization with MAB206 reduces pro-DDLPS effects of preadipocyte-derived conditioned media.** MTS evaluation of Lipo815 cells cultured in the absence or presence of PreAdip-derived CM and MAB206 (0.6 μg/mL) for 96h.

**S4 Fig. MAB206 abrogates IL6-medicated STAT activation in DDLPS.** STAT1, STAT3 and STAT5 protein expression in Lipo815 cells pretreated with MAB206 (0.6 μg/mL, 1h) prior to the addition of IL6 (10 ng/mL, 20 min).

**S5 Fig.** Illustration of *MDM2-FL* and *MDM2-ALT1* splice form mRNA.

## References

1. Ray-Coquard I, Blay JY, Italiano A, Le Cesne A, Penel N, Zhi J, et al. Effect of the MDM2 antagonist RG7112 on the p53 pathway in patients with MDM2-amplified, well-differentiated or dedifferentiated liposarcoma: An exploratory proof-of-mechanism study. Lancet Oncol [Internet]. 2012 Nov [cited 2016 Jan 20];13(11):1133–40. Available from: http://www.ncbi.nlm.nih.gov/pubmed/23084521

2. Luke JJ, D’Adamo DR, Dickson MA, Keohan ML, Carvajal RD, Maki RG, et al. The cyclin-dependent kinase inhibitor flavopiridol potentiates doxorubicin efficacy in advanced sarcomas: Preclinical investigations and results of a phase I dose-escalation clinical trial. Clin Cancer Res [Internet]. 2012 May 1 [cited 2016 Feb 16];18(9):2638–47. Available from: http://www.pubmedcentral.nih.gov/articlerender.fcgi?artid=3343204&tool=pmcentrez&rendertype=abstract

3. Steen S, Stephenson G. Current treatment of soft tissue sarcoma. Proc (Bayl Univ Med Cent) [Internet]. 2008 Oct [cited 2015 Aug 25];21(4):392–6. Available from: http://www.pubmedcentral.nih.gov/articlerender.fcgi?artid=2566912&tool=pmcentrez&rendertype=abstract

4. Henricks WH, Chu YC, Goldblum JR, Weiss SW. Dedifferentiated liposarcoma: A clinicopathological analysis of 155 cases with a proposal for an expanded definition of dedifferentiation. Am J Surg Pathol [Internet]. 1997 Mar [cited 2015 Nov 8];21(3):271–81. Available from: http://www.ncbi.nlm.nih.gov/pubmed/9060596

5. Anaya DA, Lahat G, Wang X, Xiao L, Tuvin D, Pisters PW, et al. Establishing Prognosis in Retroperitoneal Sarcoma: A New Histology-Based Paradigm. Ann Surg Oncol [Internet]. 2009 Mar 20 [cited 2019 Jan 3];16(3):667–75. Available from: http://www.springerlink.com/index/10.1245/s10434-008-0250-2

6. Toulmonde M, Bonvalot S, Méeus P, Stoeckle E, Riou O, Isambert N, et al. Retroperitoneal sarcomas: Patterns of care at diagnosis, prognostic factors and focus on main histological subtypes: A multicenter analysis of the French Sarcoma Group. Annals of oncology: official journal of the European Society for Medical Oncology / ESMO [Internet]. 2014 Mar 1 [cited 2016 Mar 1];25(3):735–42. Available from: http://annonc.oxfordjournals.org/content/25/3/735.full

7. Ballo MT, Zagars GK, Pollock RE, Benjamin RS, Feig BW, Cormier JN, et al. Retroperitoneal soft tissue sarcoma: an analysis of radiation and surgical treatment. Int J Radiat Oncol Biol Phys [Internet]. 2007 Jan 1 [cited 2016 Mar 1];67(1):158–63. Available from: http://www.sciencedirect.com/science/article/pii/S0360301606027775

8. Casadei L, Calore F, Creighton CJ, Guescini M, Batte K, Iwenofu OH, et al. Exosome-derived miR-25-3p and miR-92a-3p stimulate liposarcoma progression. Cancer Res [Internet]. 2017 Jul 15 [cited 2017 Jul 27];77(14):3846–56. Available from: http://cancerres.aacrjournals.org/content/early/2017/06/14/0008-5472.CAN-16-2984.full-text.pdf

9. Mohamed-Ali V, Goodrick S, Rawesh A, Katz DR, Miles JM, Yudkin JS, et al. Subcutaneous adipose tissue releases interleukin-6, but not tumor necrosis factor-alpha, in vivo. J Clin Endocrinol Metab [Internet]. 1997 Dec [cited 2017 Mar 29];82(12):4196–200. Available from: http://press.endocrine.org/doi/10.1210/jcem.82.12.4450

10. Carey AL, Bruce CR, Sacchetti M, Anderson MJ, Olsen DB, Saltin B, et al. Interleukin-6 and tumor necrosis factor-alpha are not increased in patients with Type 2 diabetes: evidence that plasma interleukin-6 is related to fat mass and not insulin responsiveness. Diabetologia [Internet]. 2004 Jun 28 [cited 2017 Mar 29];47(6):1029–37. Available from: http://link.springer.com/10.1007/s00125-004-1403-x

11. Fried SK, Bunkin DA, Greenberg AS. Omental and subcutaneous adipose tissues of obese subjects release interleukin-6: Depot difference and regulation by glucocorticoid. J Clin Endocrinol Metab [Internet]. 1998 Mar [cited 2017 Mar 29];83(3):847–50. Available from: http://press.endocrine.org/doi/10.1210/jcem.83.3.4660

12. Massa M, Gasparini S, Baldelli I, Scarabelli L, Santi P, Quarto R, et al. Interaction between breast cancer cells and adipose tissue cells derived from fat grafting. Aesthetic surgery journal / the American Society for Aesthetic Plastic surgery [Internet]. 2016 Mar [cited 2016 Oct 2];36(3):358–63. Available from: http://www.ncbi.nlm.nih.gov/pubmed/26499941

13. Laurent V, Guérard A, Mazerolles C, Le Gonidec S, Toulet A, Nieto L, et al. Periprostatic adipocytes act as a driving force for prostate cancer progression in obesity. Nat Commun [Internet]. 2016 Jan 12 [cited 2016 Oct 2];7:10230. Available from: http://www.nature.com/doifinder/10.1038/ncomms10230

14. Park EJ, Lee JH, Yu GY, He G, Ali SR, Holzer RG, et al. Dietary and genetic obesity promote liver inflammation and tumorigenesis by enhancing IL-6 and TNF expression. Cell [Internet]. 2010 Jan 22 [cited 2016 Oct 2];140(2):197–208. Available from: http://www.ncbi.nlm.nih.gov/pubmed/20141834

15. Park J, Euhus DM, Scherer PE. Paracrine and endocrine effects of adipose tissue on cancer development and progression. Endocr Rev [Internet]. 2011 Aug [cited 2016 Oct 2];32(4):550–70. Available from: http://www.ncbi.nlm.nih.gov/pubmed/21642230

16. Harkins JM, Moustaid-Moussa N, Chung YJ, Penner KM, Pestka JJ, North CM, et al. Expression of interleukin-6 is greater in preadipocytes than in adipocytes of 3T3-L1 cells and C57BL/6J and ob/ob mice. J Nutr [Internet]. 2004 Oct [cited 2017 Mar 29];134(10):2673–7. Available from: http://www.ncbi.nlm.nih.gov/pubmed/15465765

17. Howlett M, Menheniott TR, Judd LM, Giraud AS. Cytokine signalling via gp130 in gastric cancer. Biochimica et Biophysica Acta (BBA) - Molecular Cell Research [Internet]. 2009 [cited 2017 Apr 1];1793(11):1623–33. Available from: http://www.sciencedirect.com/science/article/pii/S0167488909001906

18. Echevarria FD, Rickman AE, Sappington RM. Interleukin-6: A constitutive modulator of glycoprotein 130, neuroinflammatory and cell survival signaling in retina. J Clin Cell Immunol [Internet]. 2016 [cited 2017 Apr 1];7(4):1–3. Available from: http://www.omicsonline.org/open-access/interleukin6-a-constitutive-modulator-of-glycoprotein-130-neuroinflammatory-and-cell-survival-signaling-in-retina-2155-9899-1000439.php?aid=77842

19. Gerhartz C, Heesel B, Sasse J, Hemmann U, Landgraf C, Schneider-Mergener J, et al. Differential activation of acute phase response factor/STAT3 and STAT1 via the cytoplasmic domain of the interleukin 6 signal transducer gp130. I. Definition of a novel phosphotyrosine motif mediating STAT1 activation. J Biol Chem [Internet]. 1996 May 31 [cited 2018 Mar 12];271(22):12991–8. Available from: http://www.ncbi.nlm.nih.gov/pubmed/8662591

20. Park J, Tadlock L, Gores GJ, Patel T. Inhibition of interleukin 6-mediated mitogen-activated protein kinase activation attenuates growth of a cholangiocarcinoma cell line. Hepatology [Internet]. 1999 Nov [cited 2017 Apr 2];30(5):1128–33. Available from: http://doi.wiley.com/10.1002/hep.510300522

21. Hideshima T, Nakamura N, Chauhan D, Anderson KC. Biologic sequelae of interleukin-6 induced PI3-K/Akt signaling in multiple myeloma. Oncogene [Internet]. 2001 Sep 20 [cited 2017 Apr 2];20(42):5991–6000. Available from: http://www.nature.com/doifinder/10.1038/sj.onc.1204833

22. Schuett H, Oestreich R, Waetzig GH, Annema W, Luchtefeld M, Hillmer A, et al. Transsignaling of interleukin-6 crucially contributes to atherosclerosis in mice. Arterioscler Thromb Vasc Biol [Internet]. 2012 Feb 1 [cited 2017 Mar 30];32(2):281–90. Available from: http://www.ncbi.nlm.nih.gov/pubmed/22075248

23. Catar R, Witowski J, Zhu N, Lücht C, Derrac Soria A, Uceda Fernandez J, et al. IL-6 trans-signaling links inflammation with angiogenesis in the peritoneal membrane. J Am Soc Nephrol [Internet]. 2016 Nov 11 [cited 2017 Mar 30];ASN.2015101169. Available from: http://www.ncbi.nlm.nih.gov/pubmed/27837150

24. Xu S, Grande F, Garofalo A, Neamati N. Discovery of a novel orally active small-molecule gp130 inhibitor for the treatment of ovarian cancer. Mol Cancer Ther [Internet]. 2013 Jun [cited 2016 Oct 2];12(6):937–49. Available from: http://www.ncbi.nlm.nih.gov/pubmed/23536726

25. Gambella A, Bertero L, Rondon-Lagos M, Verdun Di Cantogno L, Rangel N, Pitino C, et al. FISH Diagnostic Assessment of MDM2 Amplification in Liposarcoma: Potential Pitfalls and Troubleshooting Recommendations. Int J Mol Sci. 2023 Jan 10;24(2).

26. Chandler DS, Singh RK, Caldwell LC, Bitler JL, Lozano G. Genotoxic Stress Induces Coordinately Regulated Alternative Splicing of the p53 Modulators MDM2 and MDM4. Cancer Res [Internet]. 2006 Oct 1 [cited 2019 Mar 27];66(19):9502–8. Available from: http://www.ncbi.nlm.nih.gov/pubmed/17018606

27. Dias CS, Liu Y, Yau A, Westrick L, Evans SC. Regulation of hdm2 by Stress-Induced hdm2alt1 in Tumor and Nontumorigenic Cell Lines Correlating with p53 Stability. Cancer Res [Internet]. 2006 Oct 1 [cited 2019 May 1];66(19):9467–73. Available from: http://www.ncbi.nlm.nih.gov/pubmed/17018602

28. Jacob AG, Singh RK, Comiskey DF, Rouhier MF, Mohammad F, Bebee TW, et al. Stress-Induced Alternative Splice Forms of MDM2 and MDMX Modulate the p53-Pathway in Distinct Ways. Alonso MM, editor. PLoS One [Internet]. 2014 Aug 8 [cited 2019 Mar 27];9(8):e104444. Available from: http://www.ncbi.nlm.nih.gov/pubmed/25105592

29. Peng T, Zhang P, Liu J, Nguyen T, Bolshakov S, Belousov R, et al. An experimental model for the study of well-differentiated and dedifferentiated liposarcoma; deregulation of targetable tyrosine kinase receptors. Lab Invest [Internet]. 2011 Mar [cited 2018 May 2];91(3):392–403. Available from: http://www.ncbi.nlm.nih.gov/pubmed/21060307

30. Bill KLJ, Garnett J, Meaux I, Ma X, Creighton CJ, Bolshakov S, et al. SAR405838: A novel and potent inhibitor of the MDM2:p53 axis for the treatment of dedifferentiated liposarcoma. Clin Cancer Res [Internet]. 2016 Mar 1 [cited 2018 May 2];22(5):1150–60. Available from: http://www.ncbi.nlm.nih.gov/pubmed/26475335

31. Sigalas I, Calvert AH, Anderson JJ, Neal DE, Lunec J. Alternatively spliced mdm2 transcripts with loss of p53 binding domain sequences: Transforming ability and frequent detection in human cancer. Nat Med. 1996;2(8).

32. Grivennikov S, Karin E, Terzic J, Mucida D, Yu GY, Vallabhapurapu S, et al. IL-6 and Stat3 are required for survival of intestinal epithelial cells and development of colitis-associated cancer. Cancer Cell [Internet]. 2009 Feb 3 [cited 2017 Apr 2];15(2):103–13. Available from: http://www.ncbi.nlm.nih.gov/pubmed/19185845

33. Sriuranpong V, Park JI, Amornphimoltham P, Patel V, Nelkin BD, Gutkind JS. Epidermal growth factor receptor-independent constitutive activation of STAT3 in head and neck squamous cell carcinoma is mediated by the autocrine/paracrine stimulation of the interleukin 6/gp130 cytokine system. Cancer Res [Internet]. 2003 [cited 2017 Apr 2];63(11). Available from: http://cancerres.aacrjournals.org/content/63/11/2948.long

34. Wu X, Xiao H, Wang R, Liu L, Li C, Lin J. Persistent GP130/STAT3 signaling contributes to the resistance of doxorubicin, cisplatin, and MEK inhibitor in human rhabdomyosarcoma cells. Curr Cancer Drug Targets [Internet]. 2016 [cited 2017 Sep 30];16(7):631–8. Available from: http://www.ncbi.nlm.nih.gov/pubmed/26373715

35. Gritsko T, Williams A, Turkson J, Kaneko S, Bowman T, Huang M, et al. Persistent activation of STAT3 signaling induces survivin gene expression and confers resistance to apoptosis in human breast cancer cells. Clinical Cancer Research [Internet]. 2006 [cited 2017 Apr 2];12(1). Available from: http://clincancerres.aacrjournals.org/content/12/1/11.long

36. Wang Y, Li L zhi, Ye L, Niu X long, Liu X, Zhu Y qin, et al. [Chemotherapy resistance induced by interleukin-6 in ovarian cancer cells and its signal transduction pathways]. Zhonghua Fu Chan Ke Za Zhi. 2010;45(9).

37. Yu H, Yue X, Zhao Y, Li X, Wu L, Zhang C, et al. LIF negatively regulates tumour-suppressor p53 through Stat3/ID1/MDM2 in colorectal cancers. Nat Commun [Internet]. 2014 [cited 2016 Oct 2];5:5218. Available from: http://www.ncbi.nlm.nih.gov/pubmed/25323535

38. Grande F, Aiello F, Grazia O De, Brizzi A, Garofalo A, Neamati N. Synthesis and antitumor activities of a series of novel quinoxalinhydrazides. Bioorg Med Chem [Internet]. 2007 Jan 1 [cited 2018 Feb 26];15(1):288–94. Available from: https://www.sciencedirect.com/science/article/pii/S0968089606008005?via%3Dihub

39. Gu L, Findley HW, Zhou M. MDM2 induces NF-κB/p65 expression transcriptionally through Sp1-binding sites: a novel, p53-independent role of MDM2 in doxorubicin resistance in acute lymphoblastic leukemia. Blood [Internet]. 2002 [cited 2017 Apr 2];99(9). Available from: http://www.bloodjournal.org/content/99/9/3367.long?sso-checked=true

40. Zhou M, Gu L, Findley HW, Jiang R, Woods WG. PTEN reverses MDM2-mediated chemotherapy resistance by interacting with p53 in acute lymphoblastic leukemia cells. Cancer Res [Internet]. 2003 [cited 2017 Apr 2];63(19). Available from: http://cancerres.aacrjournals.org/content/63/19/6357.long

41. Koster R, Timmer-Bosscha H, Bischoff R, Gietema JA, de Jong S. Disruption of the MDM2–p53 interaction strongly potentiates p53-dependent apoptosis in cisplatin-resistant human testicular carcinoma cells via the Fas/FasL pathway. Cell Death Dis [Internet]. 2011 Apr [cited 2017 Apr 2];2(4):e148. Available from: http://www.nature.com/doifinder/10.1038/cddis.2011.33

42. Italiano A, Toulmonde M, Cioffi A, Penel N, Isambert N, Bompas E, et al. Advanced well-differentiated/dedifferentiated liposarcomas: role of chemotherapy and survival. Annals of oncology: official journal of the European Society for Medical Oncology / ESMO [Internet]. 2012 Jun 1 [cited 2015 Dec 10];23(6):1601–7. Available from: https://academic.oup.com/annonc/article-lookup/doi/10.1093/annonc/mdr485

43. Santoro A, Tursz T, Mouridsen H, Verweij J, Steward W, Somers R, et al. Doxorubicin versus CYVADIC versus doxorubicin plus ifosfamide in first-line treatment of advanced soft tissue sarcomas: a randomized study of the European Organization for Research and Treatment of Cancer Soft Tissue and Bone Sarcoma Group. J Clin Oncol [Internet]. 1995 Jul 21 [cited 2017 Apr 3];13(7):1537–45. Available from: http://www.ncbi.nlm.nih.gov/pubmed/7602342

44. Xiao H, Bid HK, Chen X, Wu X, Wei J, Bian Y, et al. Repositioning Bazedoxifene as a novel IL-6/GP130 signaling antagonist for human rhabdomyosarcoma therapy. PLoS One [Internet]. 2017 [cited 2018 May 2];12(7):e0180297. Available from: http://www.ncbi.nlm.nih.gov/pubmed/28672024

45. Evans SC, Viswanathan M, Grier JD, Narayana M, El-Naggar AK, Lozano G. An alternatively spliced HDM2 product increases p53 activity by inhibiting HDM2. Oncogene [Internet]. 2001 Jul 11 [cited 2019 Mar 27];20(30):4041–9. Available from: http://www.ncbi.nlm.nih.gov/pubmed/11494132

46. Eiro N, Fernandez-Gomez J, Sacristán R, Fernandez-Garcia B, Lobo B, Gonzalez-Suarez J, et al. Stromal factors involved in human prostate cancer development, progression and castration resistance. 2017 Feb 27 [cited 2018 Jul 17];143(2):351–9. Available from: http://link.springer.com/10.1007/s00432-016-2284-3

47. Lee J, Hong BS, Ryu HS, Lee HB, Lee M, Park IA, et al. Transition into inflammatory cancer-associated adipocytes in breast cancer microenvironment requires microRNA regulatory mechanism. PLoS One [Internet]. 2017 [cited 2018 Sep 16];12(3):e0174126. Available from: http://www.ncbi.nlm.nih.gov/pubmed/28333977

